# DR3E-Net: Drug Repurposing Network Framework Based on Deep Embedding, Differentially Expressed Genes, and Drug Side Effects

**DOI:** 10.1101/2024.06.15.599138

**Authors:** YoungMin Bhak, Joon Ho Park, Hyun Wook Han

## Abstract

In contrast to traditional de-novo drug discovery, which involves high costs and lengthy development times alongside a high failure rate, drug repurposing (DR) offers a potential solution to these challenges. This study introduces DR3E-Net, an in-silico DR framework that integrates embedded vectors of genetic expression influenced by drugs, differentially expressed genes (DEGs) associated with diseases, and information about drug side effects. The DR3E-Net utilized LINCS L1000 datasets, which comprised 680,909 samples, 3,820 drugs, and 977 genes after preprocessing. The deep-embedding model within DR3E-Net was trained on this dataset, facilitating the transformation of 977-dimensional vectors into 32-dimensional unit-hypersphere embeddings. Subsequently, cosine similarity (CS) was computed among them (Step 1). To further elaborate, agonistic drugs against DEG linked to a disease (Alzheimer’s disease (AD) in this study) were merged with outcomes from Step 1 (Step 2). Finally, the SIDER and SAEDR/DRIP databases were combined with the integrated results from Step 2 (Step 3).

The DR3E-Net was utilized to predict potential alternative drug candidates for three AD medications: donepezil, galantamine, and memantine. The drugs predicted by the deep-embedding model (Step 1) were combined with agonistic drugs against CRTAP gene, resulting in 111, 86, and 96 drugs, respectively (Step 2). Lastly, drug side effect information was incorporated with the aforementioned drug outcomes, yielding 21, 15, and 18 drugs (Step 3). DR3E-Net was developed to predict functionally similar drugs while considering disease-associated DEGs of patients and drug side effect information.

## INTRODUCTION

The identification of new therapeutic indications for existing drugs, known as drug repurposing, has gained significant attention in recent years due to its potential to expedite drug discovery processes and reduce costs.^1^ Traditional de-novo drug discovery methods often involve high-throughput screening and target-based approaches, which are time-consuming and resource-intensive.^2^ In contrast, drug repurposing capitalizes on existing knowledge about the safety profiles and mechanisms of action of approved drugs, allowing for the exploration of new therapeutic opportunities.^3^ A key challenge in drug repurposing is the identification of novel indications based on the intricate relationships between drug compounds and disease pathways. To address this challenge, computational approaches have emerged as valuable tools for efficiently sifting through large-scale omics data and drug databases. Among these, deep learning has shown promise in capturing complex patterns within large data and revealing subtle relationships using multiple processing layers and the backpropagation algorithm that may not be apparent through traditional methods.^4^ By learning intricate representations from raw data, deep learning models can unveil hidden connections and similarities that facilitate drug repurposing efforts.^5^ Regarding biological data, one notable resource that has facilitated the exploration of drug repurposing opportunities is the Library of Integrated Network-Based Cellular Signatures (LINCS) L1000 dataset.^6^ This dataset comprises gene expression profiles of human cells treated with various perturbagens, representing a comprehensive snapshot of cellular responses to pharmacological interventions. Importantly, LINCS L1000 data provides insights into the functional impact of perturbagens on cellular pathways, enabling a more nuanced understanding of drug effects beyond compound structural considerations.^7^ In this study, we propose a novel approach for drug repurposing that leverages deep learning techniques to embed the expression profiles of samples composed of 977 landmark genes from the LINCS L1000 dataset into 32-dimensional unit-hypersphere.^8^ The cosine similarity between two samples embedded as 32-dimensional unit-vectors are supposed to be high when the corresponding drugs exhibit similar genetic expression patterns.^8^ In this way, our method focuses on capturing functional similarity rather than structural resemblance among drugs.^8^ By prioritizing functional similarity over structural analogies, our approach addresses a critical limitation of traditional drug repurposing methods that rely solely on structural comparisons.^8–12^ Furthermore, our method integrates information from differentially expressed genes (DEGs) associated with specific diseases and incorporates side effect profiles to refine the drug repurposing process. By considering both disease-specific genetic alterations and potential adverse effects, our approach enhances the precision and safety of repurposed drug candidates. In summary, this research introduces a drug-repurposing framework that uses deep learning-based strategy that harnesses the power of the LINCS L1000 dataset to capture functional similarities between drugs based on their impact on gene expression profiles with comprehensively considering DEGs with respect to specific disease and side effect information of drugs.

## METHODS

### Dataset for training deep-embedding model

A deep-embedding model was trained using data derived from the LINCS L1000 study.^6^ The datasets were obtained via accession numbers GSE70138 and GSE92742 from the NCBI GEO web pages.^13,14^ The data labeled as level 4, which represents relative gene expression after a z-scoring procedure, was extracted from the respective dataset web pages.^6^ For data preprocessing of GSE70138 and GSE92742 (Figure 1), samples with a “canonical_smiles” column value of −666 were excluded. Only samples with “pert_type” column values of “trt_cp” were retained to exclude those not associated with treatment samples affected by compounds. Subsequently, 977 landmark genes from the dataset were selected to reduce dataset dimensionality. Gene-wise standardization 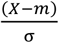 was independently applied to each dataset, followed by vertical concatenation of the two datasets. Finally, only samples with drugs occurring at least 40 times were retained, while samples with drugs starting with BRD were removed.

**Figure 1.**
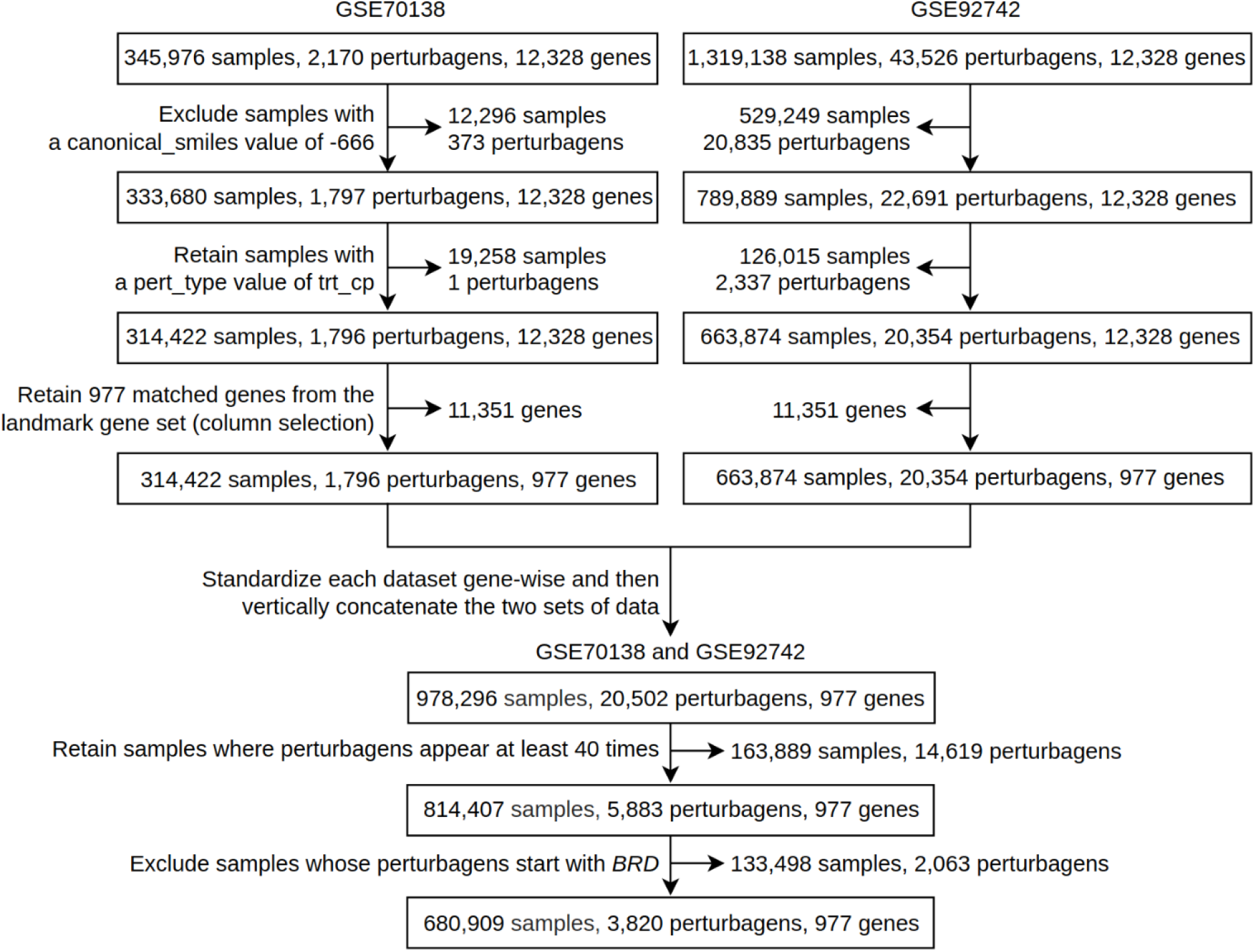
Data preprocessing for the two datasets employed in training a deep-embedding model. The identical data processing procedure was performed on two datasets (GSE70138 and GSE92742), as outlined below: 1) Samples with a “canonical_smiles” column value of −666 were removed. 2) Only samples with a “pert_type” value of “trt_cp” were retained, while others were excluded. 3) Out of the 12,328 gene columns, 977 were selected. 4) Standardization was conducted, and the outcomes were concatenated vertically. Following concatenation, the subsequent processing steps were taken: 1) Only samples with a drug frequency exceeding 40 were kept; otherwise, they were excluded. 2) Samples whose drug name starting with BRD were excluded. GSE, GEO series; trt_cp, treatment sample affected by compounds; BRD, Broad

### Overall architecture of DR3E-Net

Upon completion of training the deep-embedding model, it becomes capable of embedding the relative gene expression **x**_*src*_ ∈ ℝ^977^of entire samples into **x**_*out*_ ∈ ℝ^32^ (Figure 2, Step 1).^8^ Subsequently, a target drug was designated for the purpose of drug repurposing, and the cosine similarities (CS) were computed using dot-product of unit-vectors **x**_*drugA*_ ⋅ **x**_*drugB*_ where **x**_*drug*_ ∈ ℝ^32^ (Figure 2, Step 1).^8^ The identification of DEGs associated with a specific disease was accomplished through the compilation of information from previous research. From this, drugs that function as agonists against this disease related DEG were selected from the same LINCS L1000 data (Figure 2, Step 2). Subsequent to this, a collection of common drugs was generated by combining the predicted drugs from Step 1 with the drugs from Step 2 (Figure 2, Step 2). Lastly, the common drugs identified in Step 2 were cross-referenced with the drug side effect databases (Figure 2, Step 3).^15,16^

**Figure 2.**
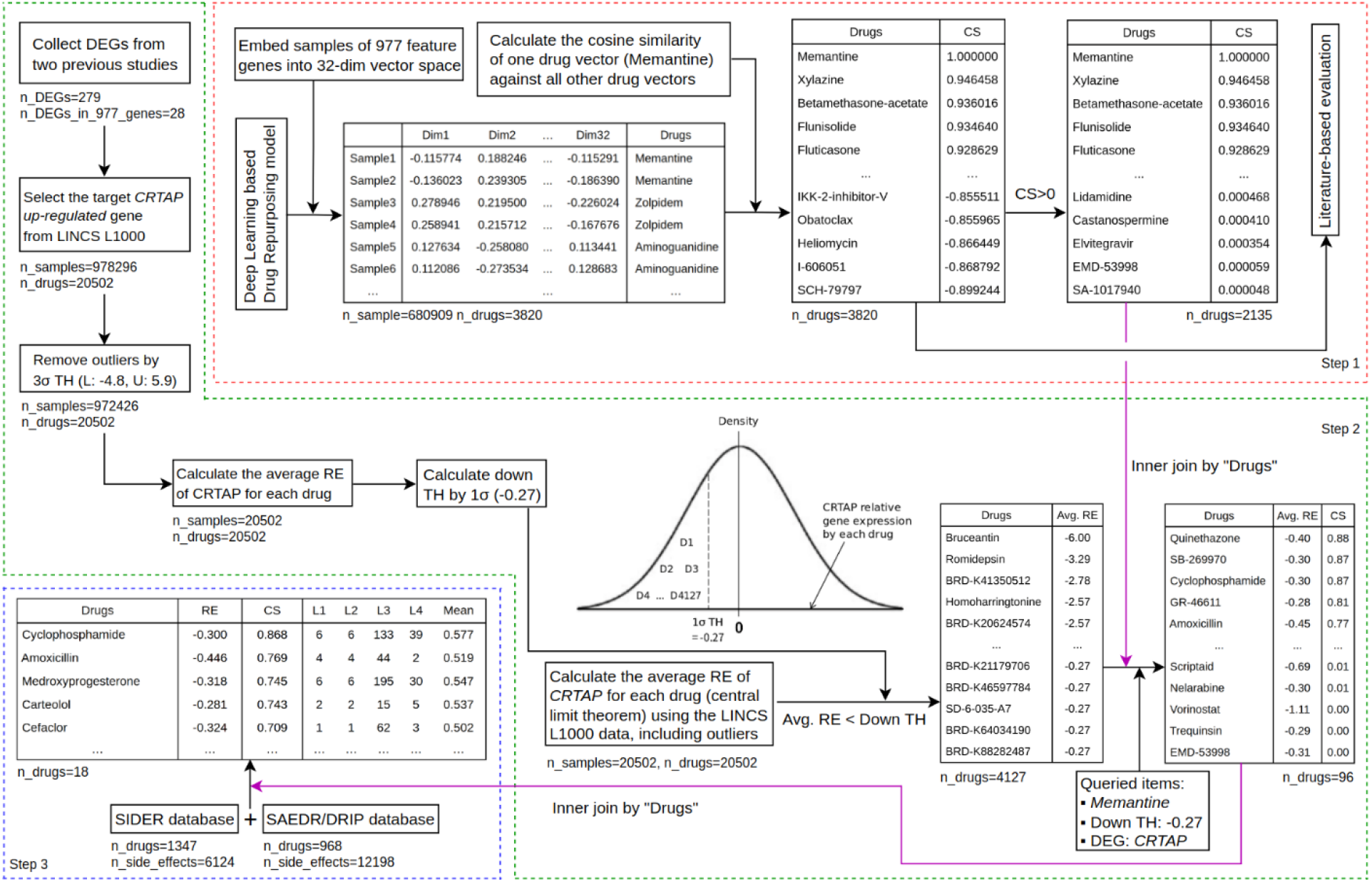
Overall architecture of DR3E-Net framework. The DR3E-Net operates through a three-step process. In the initial step, the deep-embedding model generates vectors representing samples within a 32-dimensional unit-hypersphere. Subsequently, cosine similarities are computed between a specific drug vector (memantine in the depicted figure) and all other drug vectors. Following this, a literature-based evaluation is conducted for predicted candidates from deep-embedding model. Moving to the second step, the focus is on identifying significant DEGs associated with the target disease (AD in the illustrated context). From the averaged relative expression distribution, agonistic drugs are extracted based on a down threshold computed using 1σ. The outcomes from the first and second steps are combined by the drug name. In the last stage, the concluding outcome of the second step is integrated with the SIDER and SAEDR/DRIP databases by the drug name to incorporate information about the potential side effects of drug repurposing candidates. AD, Alzheimer’s disease; CS, cosine similarity; DEGs, differentially expressed genes; L, lower bound; U, upper bound; RE, relative expression; TH, threshold; Avg, average.

### Training of deep-embedding model

The deep-embedding model was implemented through the integration of growth-rate and skip-connection structures, which were derived from DenseNet^17^ and ResNet,^18^ respectively (Figure 3). The model accommodated input source data **x**_*src*_ ∈ ℝ^977^ of 680,909 samples where 977 is the number of feature genes. These samples were fed into the deep-embedding model, producing embedded vectors **x**_*out*_ ∈ ℝ^32^. These vectors underwent L2 normalization 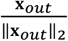 before entering the column-wise L2 normalized classifier layer 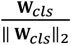, where **W**_*cls*_ ∈ ℝ^32×3820^ and 3,820 is the number of drugs.^8^ The classifier layer’s output **x**_*cls*_ ∈ ℝ^3820^, along with **x**_*out*_ ∈ ℝ^32^, drug labels **x**_*lbl*_ ∈ ℕ^680909^ (1 ≤ N ≤ 3,820) and drug weights **x**_*weight*_ ∈ ℝ^680909^, were collectively processed using an additive margin softmax loss function (Eq1), as denoted.^19–22^

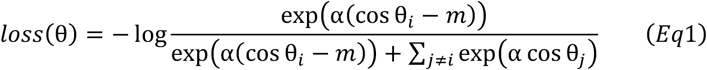

where *α* (> 0) is a trainable scaling parameter, *m*(= 0.2) is a margin hyperparameter, *i* is an index of i^th^ drug class, *j* ≠ *i* as indices of all drug classes except for i^th^ drug.^19^ And *cos* θ_*i*_ in Eq1 is represented by Eq2.^19–22^

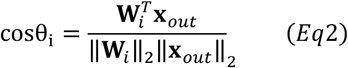

where *cos* θ_*i*_ was determined by the dot product of column-wise L2 normalized Classifier layer 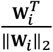 and 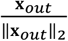.^19^ The deep-embedding model’s entire trainable parameter set ψ was updated using the ADAM optimizer^23^ guided by ∇_ψ_ ℒ (ψ, 𝒟), where 𝒟 is composed of **x**_*src*_ ∈ ℝ^977^, **x**_*lbl*_ ∈ ℕ^680909^ and **x**_*weight*_ ∈ ℝ^680909^. Hyperparameters and the development environment were explained in the Supplementary Texts.

**Figure 3.**
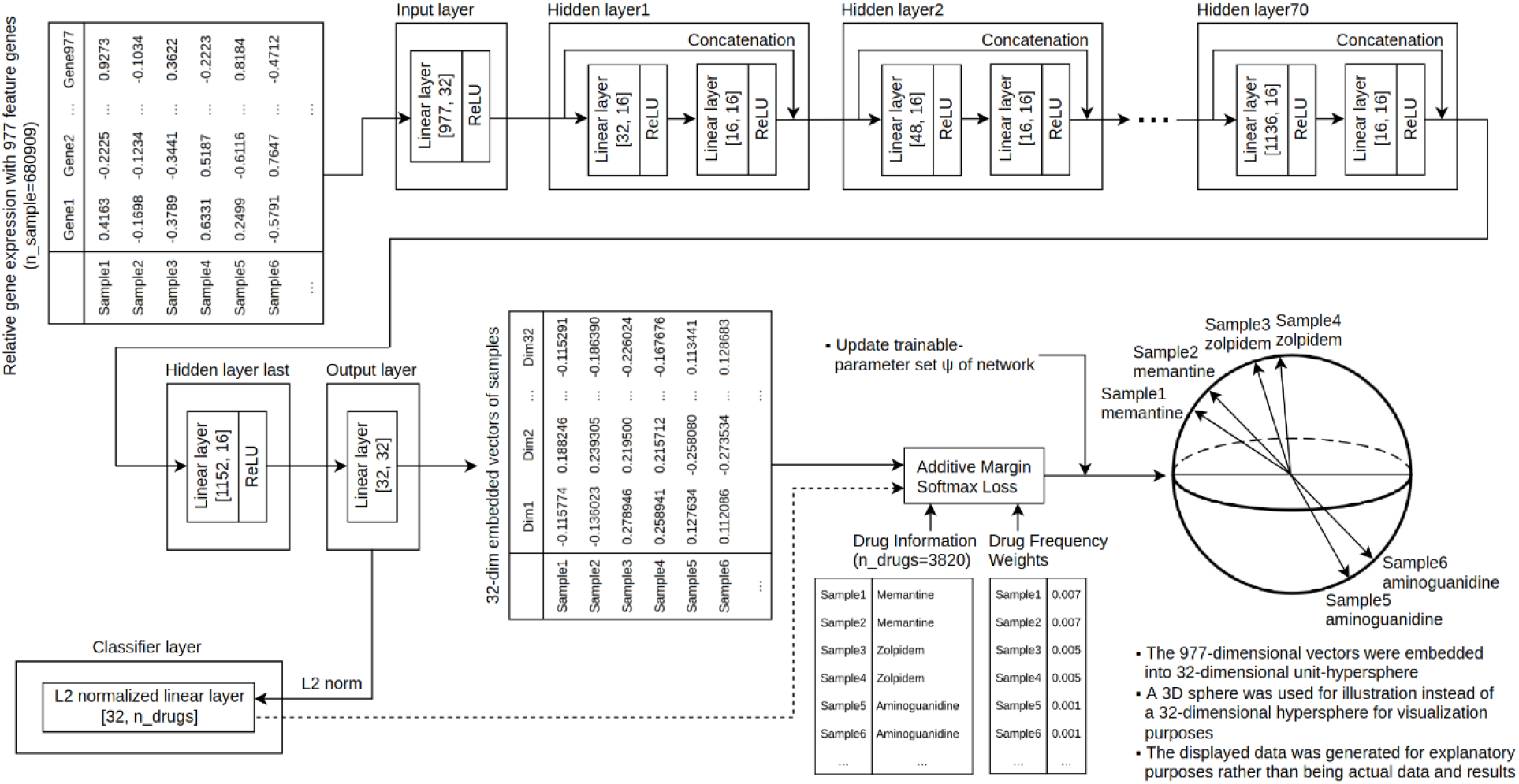
The procedure of training deep-embedding model. The deep-embedding model’s input layer receives relative gene expression samples represented by 977 gene features. The output from the input layer is then transmitted to a set of hidden layers consisting of 70 processing layers, each incorporating 16 growth-rate structures and skip-connection patterns. The resultant output from the final hidden layer is directed to the Output layer, which makes predictions of sample vectors within a 32-dimensional space. A Classifier layer, normalized using L2 normalization, takes the L2 normalized output from the Output layer. Subsequently, the outcomes from both the Classifier layer and the Output layer, in addition to drug information and drug frequency weight, are fed into an additive margin softmax loss function. The deep-embedding model’s trainable parameters are adjusted based on the gradients of the loss value. Following adequate model training, the cosine similarity value between two 32-dimensional sample vectors becomes elevated beyond the range of −1 to 1 if those were embedded with lower angle between.

### Calculating cosine similarity among drug vectors

To compute CS with respect to a specific drug, the designated drug should be indicated first.^19^ Figure 4 explains this procedure with an example of Alzheimer’s disease (AD) drug memantine as the target drug. Sample vectors **x**_*out*_ ∈ ℝ^32^ underwent L2 normalization to transform them into unit vectors 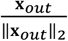.^19^ The visual representation in Figure 4 displays generated tables and vectors within a 3-dimensional sphere for explanatory and visualization purposes. Subsequently, 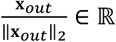^32^ were collectively averaged based on drug names, and the resulting averaged drug vectors **x**_*drug*_ ∈ ℝ^32^ also underwent L2 normalization.^19^ Following this, the drug vector corresponding to the target drug (memantine in Figure 4) was subjected to an dot product operation with all other drug vectors 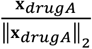⋅ 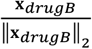, where drugB = {drug|drug, ∀_drug_} and |drugB| = 3820, facilitating the computation of CS amongst them.^19^

**Figure 4.**
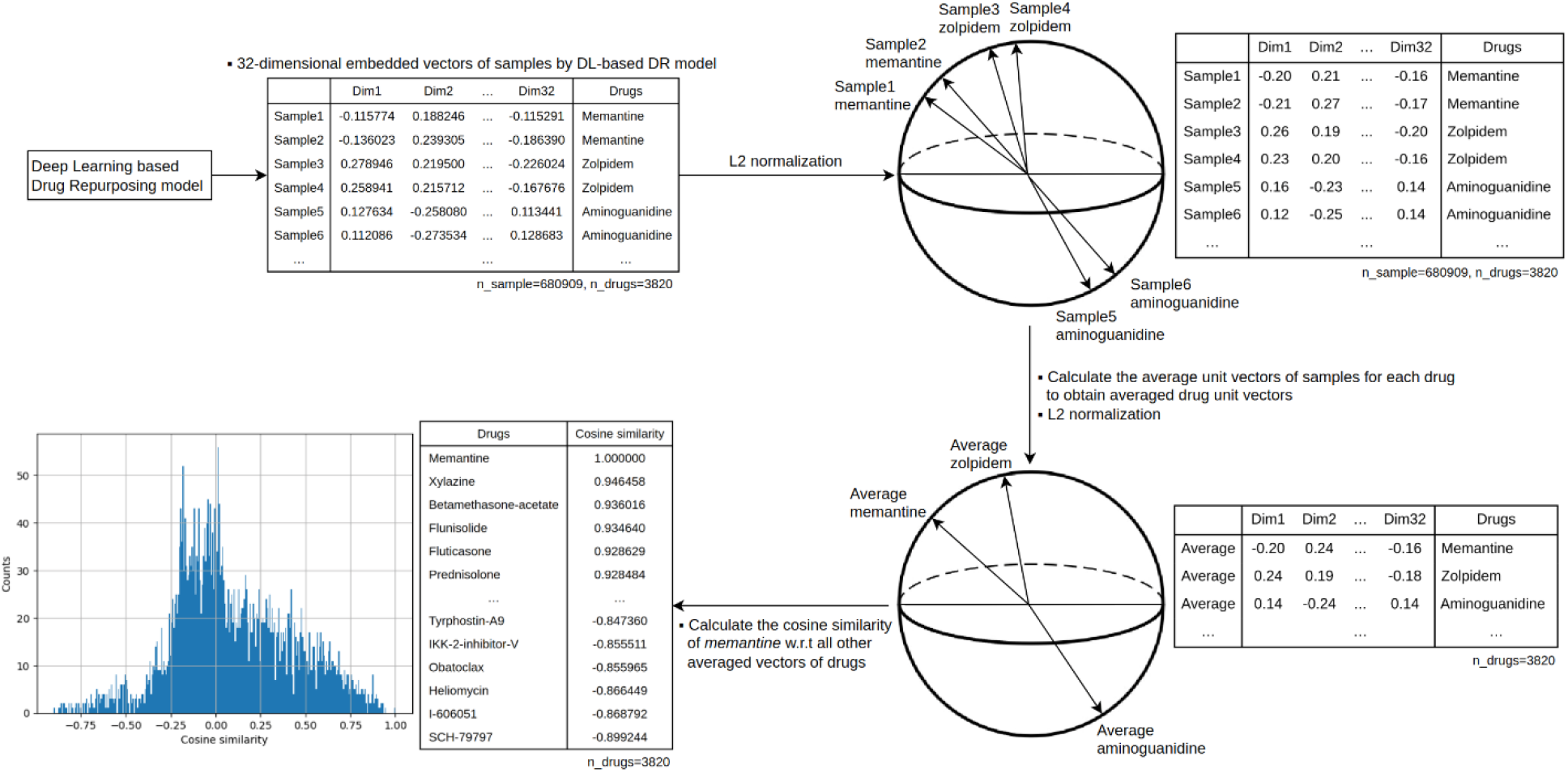
The methodology for computing cosine similarity scores. The deep-embedding model that underwent training is capable of predicting sample vectors represented within a 32-dimensional space. These vectors underwent L2 normalization to ensure their unit-vector status. Following this, the sample unit-vectors underwent averaging on a column-wise basis for each drug, followed by L2 normalization. This process yielded drug unit-vectors, each with a dimension of 32. Subsequently, a particular drug vector (memantine in the figure) was singled out, and its cosine similarity was computed against the complete set of other drug vectors.

### Literature-based evaluation of deep-embedding model

The literature-based evaluation of the deep-embedding model also took into consideration prior research.^24–35^ This model was used to predict potential drug repurposing candidates for three categories of AD medications: donepezil, galantamine, and memantine. The previously predicted candidates with significant CS from Step 1 were examined based on existing studies (Table 1).

**Table 1.** Qualitative evaluation on predicted drugs from the deep-embedding model. The deep-embedding model predicted drugs that share functional similarities in relation to three distinct categories of medications used for Alzheimer’s disease. Drugs that were projected by the deep-embedding model and exhibited high cosine similarity values were examined in earlier research investigations.

| Target: Donepezil |  |  |  |  |
| --- | --- | --- | --- | --- |
| Drugs | CS | Medical uses | Evidence study | Published journal & year |
| Febuxostat | 0.95 | To treat chronic gout and hyperuricemia | Comparative effectiveness of allopurinol versus febuxostat for preventing incident dementia in older adults: a propensity-matched analysis | Arthritis Research & Therapy (2018) |
| Miglitol | 0.95 | Anti-diabetic | Causal association evaluation of diabetes with Alzheimer's disease and genetic analysis of antidiabetic drugs against Alzheimer's disease | Cell & Bioscience (2022) |
| Hymecromone | 0.90 | Choleretic and antispasmodic drugs | Oral treatment of 4-methylumbelliferone reduced perineuronal nets and improved recognition memory in mice | Brain Research Bulletin (2022) |
| Nefiracetam | 0.87 | Nootropic drugs | Improvement by nefiracetam of β-amyloid-(1-42)-induced learning and memory impairments in rats | British Journal of Pharmacology (2009) |
| Target: Galantamine |  |  |  |  |

| Drugs | CS | Medical uses | Evidence study | Published journal & year |
| --- | --- | --- | --- | --- |
| Fursultiamine | 0.98 | To treat thiamine deficiency | Thiamine therapy in Alzheimer's disease | Metabolic Brain Disease (2005) |
| Cycloserine-(d) | 0.97 | Anti-tuberculosis drug | Unraveling Comparative Anti-Amyloidogenic Behavior of Pyrazinamide and D-Cycloserine: A Mechanistic Biophysical Insight | PLOS ONE (2015) |
| Sulfinpyrazone | 0.95 | Antigout agents | Enhancing autophagy in Alzheimer's disease through drug repositioning | Pharmacology & Therapeutics (2022) |
| Naringin | 0.94 | Flavonoid | Neuropharmaceutical Properties of Naringin Against Alzheimer's and Parkinson's Diseases | Journal of Food Biochemistry (2022) |
| **Target: Memantine** |  |  |  |  |
| Drugs | CS | Medical uses | Evidence study | Published journal & year |
| Fluticasone | 0.93 | To treat nasal symptoms | High-Throughput Clinical Trial Emulation with Real World Data and Machine Learning: A Case Study of Drug Repurposing for Alzheimer's Disease | Europe PMC (2022) |
| Prednisolone | 0.93 | To treat certain types of allergies, inflammatory conditions, autoimmune disorders, and cancers. | Methylprednisolone alleviates cognitive functions through the regulation of neuroinflammation in Alzheimer's disease | Frontiers in Immunology (2023) |
| Quipazine | 0.90 | Serotonergic drug | Alzheimer's disease master regulators analysis: search for potential molecular targets and drug repositioning candidates | Alzheimer's Research & Therapy (2018) |
| Fluocinonide | 0.89 | Anti-inflammatory agent | Drug repurposing for Alzheimer's disease based on transcriptional profiling of human iPSC-derived cortical neurons | Translational Psychiatry (2019) |

CS, cosine similarity

### Agonistic drugs against differentially expressed genes

In order to identify DEGs associated with AD, we conducted a review of two prior studies.^36,37^ Additionally, a specific up-regulated DEG was chosen based on information from a previous patent document.^38^ Subsequently, only the target DEG column was retained within the preprocessed dataset **X** ∈ ℝ^978296×977^, serving the purpose of excluding outliers using the 3σ criterion in the targeted DEG column, resulting in **x**_*DEG wo outliers*_ ∈ ℝ^972426^. Next, an averaging operation was conducted based on drug names for **x**_*DEG wo outliers*_ ∈ ℝ^972426^, resulting in **x**_*DEG wo outliers*_ ∈ ℝ^20502^. Subsequently, from **x**_*DEG wo outliers*_ ∈ ℝ^20502^, a down-regulation threshold for DEG was established using the value of 1σ. Afterwards, the relative gene expressions of the target DEG column **x**_*DEG w outliers*_ ∈ ℝ^978296^ were averaged for each drug, and drugs whose average relative expressions fell below the value of 1σ calculated from the previous step were identified as agonistic drugs against the targeted DEG. The set of identified agonistic drugs was subsequently merged with the list of projected drugs showing positive CS, which was generated by the deep-embedding model, utilizing an inner join approach based on drug names (Figure 2, Step 2).

### Considering drug side effects in predicting drug repurposing candidates

The acquisition of drug-related side effect data involved the utilization of the Side Effect Resource (SIDER) and Severity of Adverse Events Derived from Reddit (SAEDR)/DRIP (Drug Risk Profile) databases.^15,16^ The SIDER dataset consists of pairs of drugs and their associated side effects, while the SAEDR/DRIP dataset provides severity scores for drug side effects and drug risk score. Both the SIDER and SAEDR/DRIP datasets were preprocessed, resulting in **X**_*SIDER*_ ∈ ℤ^1347×6124^, where *Z* = {0,1}, 1347 and 6124 are the number of drugs and side effects, and **X**_*SAEDR*_ ∈ ℝ^12198×4^, where 12198 and 4 are the number of drug side effects and the columns representing the drug side effect codes, side effect names, severity scores, and severity categories (level 1 (mild): less than 25%, level 2: 50%, level 3: 75%, level 4 (severe): greater than 75% in severity score ranged from 0 to 1) as well as **X**_*DRIP*_ ∈ ℝ^938×2^, where 938 and 2 are the number of drugs and 2 columns indicating drug codes and drug risk scores. The integrated SIDER and SAEDR/DRIP data were merged based on drug names with the drug output obtained from Step 2 in Figure 2 (Figure 2, Step 3)

## RESULTS

### Predicted drugs from deep-embedding model

A deep-embedding model that underwent training was utilized to predict functionally similar drugs with high CS (−1 ≤ CS ≤ 1) against three existing AD medications (donepezil, galantamine, and memantine). Among a pool of 3820 drugs used for deep-embedding model, 1879, 1898, and 2135 drugs displayed positive CS values in relation to the three AD drugs. Further details, including a comprehensive list of CS values and their distribution, can be accessed in the Supplementary table sheet 1 to 3.

### Review of relevant studies to validate predicted drugs functionally similar to AD medications

For qualitative assessment, predicted drugs by deep-embedding model with high CS to three AD medications were conducted by comparing them with established findings from previous studies. Notably, febuxostat, prescribed for chronic gout and hyperuricemia, exhibited the highest CS of 0.95, indicating the strongest functional resemblance to donepezil. Similarly, fursultiamine, utilized for treating thiamine deficiency, demonstrated a remarkable CS of 0.98 in relation to galantamine. Another instance was fluticasone, employed for alleviating nasal symptoms, which displayed a significant CS of 0.93 with memantine. Additional validated potential candidates identified by the deep-embedding model per three AD drugs were tabulated in Table 1.

### Integrating agonistic drug information to in-vitro drug-repurposing procedure

A total of 279 DEGs associated with AD were gathered from two preceding research studies. Among these, 28 genes were matched with the 977 deep-embedding model’s feature genes. For a comprehensive list of the analyzed DEGs, please refer to Supplementary table sheet 4. From this set of 28 DEGs, one gene of particular interest was CRTAP, an up-regulated gene to AD. This selection was based on evidence indicating its potential as a target for diagnosing and treating AD, as indicated by patents. The 3σ values used to exclude outliers from the relative gene expression of CRTAP were −4.8 and 5.9 for the lower and upper bounds, respectively. Additionally, the down-regulation threshold of 1σ, utilized to identify agonistic drugs against up-regulated CRTAP, was −0.27.

Subsequently, a total of 4,127 drugs, out of 20,502 initially considered, were identified as agonistic drugs to the CRTAP DEG. These 4,127 drugs were cross-referenced with the drugs predicted by the deep-embedding model, focusing on the drugs donepezil, galantamine, and memantine. This comparison yielded a selection of 111 drugs out of 1,879 for donepezil, 86 drugs out of 1,898 for galantamine, and 96 drugs out of 2,135 for memantine. A full list of merged information can be found in Supplementary table sheet 5.

### Adding side effect information to in-vitro drug-repurposing procedure

The SIDER and SAEDR/DRIP databases underwent preprocessing, yielding 6,124 instances of side effect information for 1,347 distinct drugs, the 12,198 pairs of drug side effects and their severity scores, and the risk score of 935 number of drugs, respectively. After preprocessing, they were integrated with the conclusive outcomes from Step 2 in Figure 2, utilizing drug names as the basis for integration. This integration yielded 21 candidate drugs from donepezil, 15 from galantamine, and 18 from memantine. The supplementary table sheet 6 to 8 contain a comprehensive analysis of the side effects of those drugs. These candidates were repurposed for potential use against AD, taking into account both their agonistic action on CRTAP DEG and associated side effects. Table 2 displays the arrangement of the top seven candidate drugs regarding CS for AD, while a complete list of results can be found in Supplementary table sheet 9.

**Table 2.** Drug repurposing candidates for Alzheimer’s disease identified through DR3E-Net’s integration of deep-embedding based functional similarity of drugs, DEG against AD, and drug side effect information.

| Target drug: Donepezil, Target DEG: CRTAP, Down-threshold for agonistic DEG: -0.27 |  |  |  |  |  |  |  |  |  |
| --- | --- | --- | --- | --- | --- | --- | --- | --- | --- |
| Drugs | RE | CS | SE<sup>a</sup><br>(n) | SI<sup>b</sup><br>(%) | S2<sup>b</sup><br>(%) | S3<sup>b</sup><br>(%) | S4<sup>b</sup><br>(%) | SAEDR<br>(Mean) | DRIP |
| Nelarabine | -0.305 | 0.386 | 84 | 1 | 1 | 89 | 8 | 0.544 | 2.277 |
| Ixabepilone | -1.286 | 0.373 | 97 | 5 | 5 | 73 | 16 | 0.557 | 2.483 |
| Nostigmine | -0.294 | 0.284 | 35 | 3 | 3 | 86 | 9 | 0.497 | 0.121 |
| Cabazitaxel | -1.76 | 0.283 | 75 | 3 | 3 | 85 | 9 | 0.534 | 1.59 |
| Etoricoxib | -0.403 | 0.283 | 129 | 5 | 5 | 78 | 12 | 0.522 | 0.508 |
| Medroxyprogesterone | -0.318 | 0.234 | 237 | 3 | 3 | 82 | 13 | 0.547 | 1.183 |
| Streptozotocin | -0.402 | 0.229 | 31 | 3 | 3 | 90 | 3 | 0.555 | 0.129 |
| **Target drug: Galantamine, Target DEG: CRTAP, Down-threshold for agonistic DEG: -0.27** |  |  |  |  |  |  |  |  |  |
| Drugs | RE | CS | SE <sup>a</sup><br>(n) | S1 <sup>b</sup><br>(%) | S2 <sup>b</sup><br>(%) | S3 <sup>b</sup><br>(%) | S4 <sup>b</sup><br>(%) | SAEDR<br>(Mean) | DRIP |
| Streptozotocin | -0.402 | 0.889 | 31 | 3 | 3 | 90 | 3 | 0.555 | 0.129 |
| Amifostine | -0.314 | 0.749 | 36 | 14 | 14 | 50 | 22 | 0.533 | N/A |
| Cisplatin | -0.366 | 0.715 | 83 | 4 | 4 | 73 | 19 | 0.555 | 0.604 |
| Carteolol | -0.281 | 0.607 | 24 | 8 | 8 | 62 | 21 | 0.537 | N/A |
| Cyclophosphamide | -0.3 | 0.296 | 184 | 3 | 3 | 72 | 21 | 0.577 | 0.743 |
| Cefaclor | -0.324 | 0.287 | 67 | 1 | 1 | 93 | 4 | 0.502 | 0.355 |
| Carbimazole | -0.291 | 0.241 | 27 | 4 | 4 | 89 | 4 | 0.509 | N/A |
| **Target drug: Memantine, Target DEG: CRTAP, Down-threshold for agonistic DEG: -0.27** |  |  |  |  |  |  |  |  |  |
| Drugs | RE | CS | SE <sup>a</sup><br>(n) | S1 <sup>b</sup><br>(%) | S2 <sup>b</sup><br>(%) | S3 <sup>b</sup><br>(%) | S4 <sup>b</sup><br>(%) | SAEDR<br>(Mean) | DRIP |
| Cyclophosphamide | -0.3 | 0.868 | 184 | 3 | 3 | 72 | 21 | 0.577 | 0.743 |
| Amoxicillin | -0.446 | 0.769 | 54 | 7 | 7 | 81 | 4 | 0.519 | 0.203 |
| Medroxyprogesterone | -0.318 | 0.745 | 237 | 3 | 3 | 82 | 13 | 0.547 | 1.183 |
| Carteolol | -0.281 | 0.743 | 24 | 8 | 8 | 62 | 21 | 0.537 | N/A |
| Cefaclor | -0.324 | 0.709 | 67 | 1 | 1 | 93 | 4 | 0.502 | 0.355 |
| Fenofibrate | -0.288 | 0.583 | 114 | 6 | 6 | 75 | 12 | 0.519 | 0.647 |
| Mafenide | -0.346 | 0.436 | 16 | 25 | 25 | 50 | 0 | 0.432 | 0.076 |

<sup>a</sup> The number of side effects of each drug
<sup>b</sup> The percentage of side effects for each SAEDR score level was rounded up.
RE, relative gene expression; CS, cosine similarity; SE, side effects; S1, SAEDR score level 1, SAEDR, Severity of Adverse Events Derived from Reddit; DRIP, drug-risk profile

## DISCUSSION

### Considerations in preprocessing dataset

When preprocessing the datasets, samples with a low frequency of drugs were excluded since they did not provide comprehensive information to the deep learning model. Despite an earlier study indicating a 90% sensitivity in a predictive model using 20 training samples,^39^ the deep-embedding model utilized a larger sample size, exceeding 40 for each drug class. The deep-embedding model, in conjunction with the analytic scenario, did not require the test data, so making a sample size of 40 per drug class might be acceptable for model training. This quantity was also in line with the general criterion of “greater than 25 or 30”, as suggested for the central limit theorem.^40^ This parameter 40 was employed to identify agonistic medications that exhibited low relative gene expression within the CRTAP gene, derived from the mean distribution of CRTAP’s relative gene expression for each drug. An additional consideration was the exclusion of samples associated with drugs whose names began with BRD. These BRD drugs were deemed unsuitable for drug repurposing due to their lack of clinical utilization after FDA approval.^6^ Among the FDA-unapproved drugs, only those associated with BRD were excluded, as they constituted a significant portion of the overall FDA-unapproved drug category. Furthermore, reducing the number of classes during deep learning model training could lead to a decrease in model complexity.^41^

### Class imbalance and batch-size in training deep-embedding model

During the training stage, we noted a class imbalance and examined the skewed distribution of drug frequency in supplementary table sheet 10. This class imbalance could lead the model to exhibit bias towards the patterns found in high frequency drug samples.^42^ To address this issue, we employed a cost-sensitive learning approach as algorithm-level method.^43^ This approach assigns varying weights to the loss values from the majority and minority classes, thereby mitigating the impact of the class imbalance.^43^ And, to optimize the training process, we determined that a mini batch-size of 700 was suitable after empirical testing, as there were indications that a large mini batch-size could be detrimental to the model training process.^44^

### The effect of normalization methods to the distribution of cosine similarity

Two techniques were experimented with data normalization: min-max normalization and standardization. While empirically min-max normalization led to a non-bell-shaped distribution of CS, the application of standardization resulted in a bell-shaped distribution. This bell-shaped pattern, achieved by standardization, is deemed suitable and optimal for the task at hand. It ensures that highly similar or dissimilar functional drugs to the target drug are sparse, while drugs with limited functional association are more frequent.

### Agonistic drugs targeting multiple DEGs in drug repurposing

The investigation focused on identifying agonistic drugs for the solitary up-regulated CRTAP DEG pattern in the aforementioned example of drug repurposing for AD. However, DR3E-Net is not restricted to utilizing only a single DEG; it has the capability to reflect multiple DEG patterns. To illustrate, suppose the DEGs of interest encompass both an up-regulated DEG1 and a down-regulated DEG2 associated with a specific disease. In this scenario, agonistic drugs that simultaneously down-regulate DEG1 and up-regulate DEG2, like the method described earlier, can be identified. Consequently, DR3E-Net exhibits the capacity to target multiple DEGs concurrently. We suggested scenario based practical application of DR3E-net for personalized and effective drug repurposing in the Supplementary Texts.

### LIMITATIONS

This investigation relies on the LINCS L1000 dataset, rendering it susceptible to potential disruptions caused by noise and measurement errors. Moreover, when contemplating DEGs, additional viewpoints, such as disease pathways and gene ontology structures, could be incorporated to identify more noteworthy target DEGs.

Additionally, the available side effect data sourced from the SIDER database encompasses a limited assortment of drugs, resulting in the reduced number of repurposing candidates within the final analytical findings. Lastly, it is important to note that this approach lacks validation through in-vivo or animal experimentation, underscoring the necessity for further validation to assess the safety of drug repurposing candidates predicted through in-vitro methodologies.

## Acknowledgement

This research was supported by the Bio Industry Technology Development Program (No. 20015086) funded by the Ministry of Trade, Industry, & Energy (MOTIE, Korea) and by the National Research Foundation of Korea (NRF) grant funded by the Korea government (MSIT)(NRF-2019M3C7A1032262).

## Data availability

GSE70138 avaliable at: https://www.ncbi.nlm.nih.gov/geo/query/acc.cgi?acc=GSE70138

GSE92742 avaliable at: https://www.ncbi.nlm.nih.gov/geo/query/acc.cgi?acc=GSE92742

## Software availability

The source code of DR3E-Net is available at https://github.com/ymb943/DR3E-Net

## Supplementary Information

DR3E-Net_supplementary_tables - xlsx file DR3E-Net_supplementary_texts - docx file

## Author contributions

YoungMin Bhak - Conceptualization, Methodology, Software, Formal analysis, Data Curation, Writing (Original Draft), Writing (Review & Editing), Visualization; Joon Ho Park - Investigation, Supervision; Hyun Wook Han - Conceptualization, Investigation, Supervision, Project administration, Funding acquisition

