## Supplementary texts for "DR3E-Net: Drug Repurposing Network Framework Based on Deep Embedding, Differentially Expressed Genes, and Drug Side Effects"

### Hyperparameters and the development environment for the model training

Hyperparameters and the development environment were configured as follows: mini-batch size of 700 with shuffling, growth-rate set to 16, 70 layers, a learning rate (lr) of 0.001 for the model, lr of 1.0 for the scaling parameter, weight decay values of 0 for both model and scaling parameter, a lr scheduler $\gamma$ of 0.5 with a decrease every 10 epochs, and a total of 80 training epochs. The model training was conducted on a Linux Ubuntu 20.04 server equipped with an Intel(R) Xeon(R) Gold 5218R CPU @ 2.10GHz, boasting 80 processors, 503GB of RAM, and a single GeForce RTX 3090 Graphics Processing Unit (GPU). The implementation utilized PyTorch 1.11.0 [1], Python 3.7 [2], Cudatoolkit 11.3[3] and libraries such as NumPy 1.21.6[4] Pandas 1.3.5 [5].

#### How to practically apply DR3E-Net framework for predictive, preventive, and personalized drug repurposing

We present a potential scenario demonstrating how the DR3E-Net framework can be harnessed to achieve an effective and personalized DR. To illustrate, let us consider an instance where a patient with AD, referred to as patient A, exhibits an up-regulation of DEG1 and a down-regulation of DEG2. Furthermore, this patient experiences side effect B from current AD medications. In this context, the deep-embedding model becomes instrumental in identifying drugs that display functional similarities to the existing AD treatments. As previously highlighted, the DR3E-Net framework facilitates the discovery of drugs acting as agonists against specific DEGs. Once drugs that exhibit functional similarity to the existing AD drug and agonistically impact DEG1 and DEG2 are identified, the resulting list of drugs can be combined with the integrated SIDER and SAEDR/DRIP database or any other sources of drug side effect information. Subsequently, the final selection of repurposed drug candidates for patient A can be made by isolating drugs that do not induce side effect B and do not have high side effect severity and drug risk score. This approach enables the DR3E-Net framework to effectively pinpoint alternative drug candidates for AD that exert agonistic effects on DEG1 and DEG2 while avoiding the occurrence of side effect B regarding patient A.
